# Rapid inactivation of SARS-CoV-2 on copper touch surfaces determined using a cell culture infectivity assay

**DOI:** 10.1101/2021.01.02.424974

**Authors:** Catherine Bryant, Sandra A. Wilks, C. William Keevil

## Abstract

COVID-19, caused by SARS-CoV-2, was first reported in China in 2019 and has transmitted rapidly around the world, currently responsible for 83 million reported cases and over 1.8 million deaths. The mode of transmission is believed principally to be airborne exposure to respiratory droplets from symptomatic and asymptomatic patients but there is also a risk of the droplets contaminating fomites such as touch surfaces including door handles, stair rails etc, leading to hand pick up and transfer to eyes, nose and mouth. We have previously shown that human coronavirus 229E survives for more than 5 days on inanimate surfaces and another laboratory reproduced this for SARS-CoV-2 this year. However, we showed rapid inactivation of Hu-CoV-229E within 10 minutes on different copper surfaces while the other laboratory indicated this took 4 hours for SARS-CoV-2. So why the difference? We have repeated our work with SARS-CoV-2 and can confirm that this coronavirus can be inactivated on copper surfaces in as little as 1 minute. We discuss why the 4 hour result may be technically flawed.

## INTRODUCTION

COVID-19, a severe acute respiratory disease caused by coronavirus SARS-CoV-2, was first reported in China in December 2019^1^ and has since transmitted rapidly around the world, currently responsible for 83 million reported cases and over 1.8 million deaths^2^. The mode of transmission is believed principally to be airborne exposure to respiratory droplets from symptomatic and asymptomatic patients produced during talking, coughing and sneezing but there is also a risk of the droplets contaminating fomites such as touch surfaces including door handles, push plates, stair rails, computer screens etc, leading to hand pick up and transfer to eyes, nose and mouth^3^. We have previously shown that human coronavirus 229E (Hu-CoV-229E), a cause of the common cold, survives for more than 5 days on inanimate surfaces such as metals, plastics and glass^4^, and another laboratory reproduced this for SARS-CoV-2 this year^5^. Moreover, we showed rapid inactivation of Hu-CoV-229E within 10 minutes on different copper surfaces

The biocidal properties of copper have been known for centuries. Contaminated surfaces are known to contribute to infection spread, therefore the use of bactericidal and anti-viral surfaces could potentially reduce the incidence of horizontal disease. The potential use of copper alloys as microbiocidal surfaces has demonstrated excellent antibacterial and antiviral activity against a range of pathogens in laboratory studies^6,7^. Copper ion release has been found to be essential to maintaining efficacy, but the mechanism of action is dependent on whether pathogens are Gram-negative or Gram-positive bacteria and enveloped or nonenveloped viruses4,7,8 (4,7,8). Significant reductions in microbial bioburden and acquisition of nosocomial infection have also been observed in clinical trials of incorporation of copper alloys in health care facilities (9, 10).

While copper appears to be a potent antibacterial and antiviral metal, causing rapid inactivation of pathogens on touch surfaces in just minutes, why did van Doremalen *et al*. (5) indicate that copper took 4 hours to inactivate SARS-CoV-2? To answer this, we have repeated our work with SARS-CoV-2 and can confirm that this coronavirus can indeed be inactivated on pure copper and copper-coated surfaces in as little as 1 minute. We discuss why the 4-hour inactivation result may be technically flawed due to how the virus is propagated.

## MATERIALS & METHODS

### Viral strains and cell lines

Human coronavirus SARS-CoV-2 and a kidney cell line, VERO-E6, were supplied by Public Health England (PHE), UK. Cells were maintained in minimal essential medium (MEM) supplemented with GlutaMax-1, nonessential amino acids, and 5% foetal calf serum and incubated at 37°C and 5% CO_2_. Cells were passaged twice a week using trypsin (0.25%)-EDTA and were not used beyond passage 30 (P30) (which occurred before the onset of senescence, but susceptibility to infection diminished greatly from P30). Viral stocks were prepared by infecting cells at multiplicity of infection of 0.01 for 4 to 7 days until a significant cytopathic effect (CPE) was observed. Infected cell supernatant was stored at −80°C.

### Preparation of control sample surfaces

The metal coupons (10 x 10 x 0.5 mm) were degreased in acetone, stored in absolute ethanol, and flamed prior to use. Copper (C11000) and stainless steel (S30400) coupons were supplied by the Copper Development Association. Copper-coated stainless steel coupons, produced by spraying pure copper powder cold at high pressure to form a permanent bond with the base metal to 150 micron thickness, were supplied by Copper Cover Ltd (Annahagh, Ireland).

### Infectivity assay for SARS-CoV-2 exposed to solid surfaces

Infected cell supernatant preparations of SARS-CoV-2 were spread over the metal coupons, either 1 μl to simulate hand transfer or 10 μl to simulate respiratory droplet contact, and incubated at room temperature. The virus was removed from the test surfaces after various time points using infection medium and assayed for infectious virus survival by a plaque assay, quantified as plaque forming units (PFU). Briefly, dilutions were prepared in infection medium, and 400 μl aliquots were plated onto confluent monolayers of VERO-E6 cells that had been prepared 24 h earlier in 12-well plates. The inoculum was removed after 60 min and replaced with Avicel overlays, and plates were incubated at 37°C and 5% CO_2_ for 3 days. The monolayers were fixed for 30 minutes in 8% (w/v) paraformaldehyde, stained with crystal violet, allowed to dry and plaques in the monolayer enumerated. An example of a plaque assay is shown in Figure 1.

**Figure 1.**
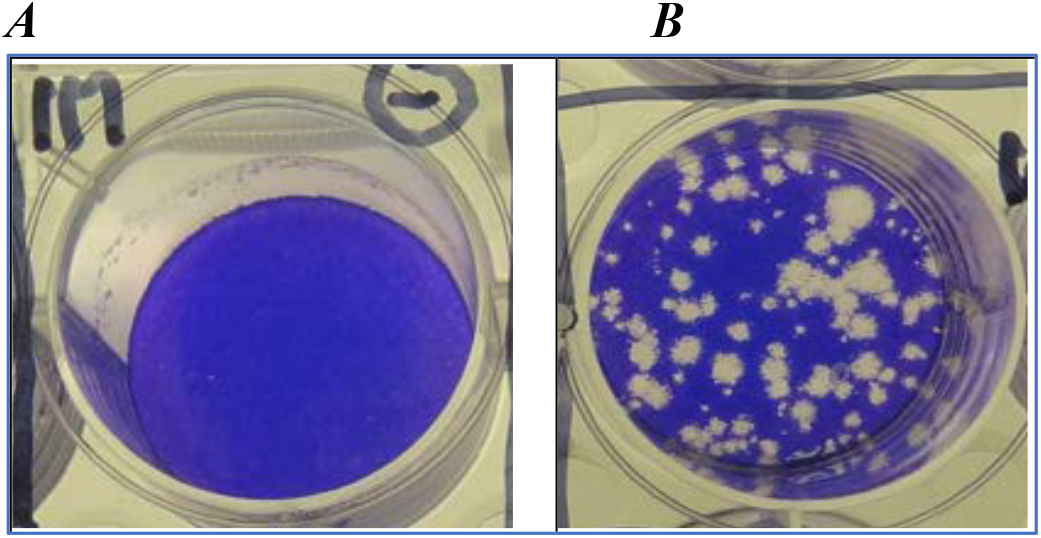
Plaque Assay with SARS-CoV-2. A) uninfected VERO-E6 cells grown to confluency and stained with crystal violet to show take up by viable cells. B) confluent VERO-E6 cells infected with SARS-CoV-2 and subsequently stained with crystal violet to show dead and dying cells seen as clear plaques.

**Table 1.** SARS-CoV-2 survival as a 1 μl inoculum to simulate hand transfer on polyethylene, stainless steel, copper coated stainless steel and pure copper coupons. Data show plaque forming units recovered from each surface whilst those in parenthesis show log decrease (n=3).

| **Dry** | **1 min** | **10 min** | **30 min** | **1 h** | **3 h** | **24 h** |
| --- | --- | --- | --- | --- | --- | --- |
| Polyethylene | 550 | 550 | 550 | 550 | 550 | 550 |
| SS | 550 | 550 | 550 | 550 | 550 | 550 |
| Cu coated SS | 0<br>(2.74) | 0<br>(2.74) | 0<br>(2.74) | 0<br>(2.74) | 0<br>(2.74) | 0<br>(2.74) |
| Cu | 0<br>(2.74) | 0<br>(2.74) | 0<br>(2.74) | 0<br>(2.74) | 0<br>(2.74) | 0<br>(2.74) |

**Table 2.** SARS-CoV-2 survival as a 10 μl inoculum to simulate respiratory droplet contact on polyethylene, stainless steel, copper coated stainless steel and pure copper coupons. Data show plaque forming units recovered from each surface whilst those in parenthesis show log decrease (n=3).

| **Moist** | **1 min** | **10 min** | **30 min** | **1 h** | **3 h** | **24 h** |
| --- | --- | --- | --- | --- | --- | --- |
| Polyethylene | 5500 | 5500 | 5500 | 5500 | 5500 | 5500 |
| SS | 5500 | 5500 | 5500 | 5500 | 5500 | 5500 |
| Cu coated SS | 5500 | 0<br>(3.74) | 0<br>(3.74) | 0<br>(3.74) | 0<br>(3.74) | 0<br>(3.74) |
| Cu | 5500 | 0<br>(3.74) | 0<br>(3.74) | 8<br>(2.83) | 0<br>(3.74) | 0<br>(3.74) |

## RESULTS

When SARS-CoV-2 was spread over the metal coupons using a 1 μl inoculum to simulate hand transfer, this dried in seconds. The initial challenge of 550 PFU was not reduced when recovered from stainless steel or polyethylene after 24 hours contact at room temperature. By contrast, no viable virus could be recovered from the copper or copper-coated stainless steel after 1 minute exposure, equivalent to >log 2.74 inactivation.

Using a 10 μl inoculum to simulate respiratory droplet contact, this took over 10 minutes to dry when incubated at room temperature. This initial challenge of 5500 PFU was also not reduced when recovered from the stainless steel or polyethylene surfaces after 24 hours contact at room temperature. Now, however, there was no viral inactivation on the copper and copper-coated stainless steel surfaces after 1 minute exposure but complete inactivation occurred within 10 minutes, equivalent to >log 3.74 reduction.

## DISCUSSION

The study by van Doremalen et al. (5) was important in providing the first evidence that the new SARS-CoV-2 was capable of surviving many days on inanimate surfaces, posing a potential threat of indirect transmission when these surfaces were not cleaned or lacked antiviral activity. We has also shown previously survival of the similar HuCoV-229E on plastics, ceramics, stainless steel and glass for 4-5 days (4), as did Duan et al. for SARS-CoV-1 (11). Importantly, we showed HuCoV-229E was inactivated on copper in just minutes and its RNA destroyed (5). However, van Doremalen *et al*. reported survival on copper for 4 hours, while ours took just minutes (4). Why? One possible explanation is that they used a different virus cultured in VERO-E6 kidney cells while we cultured HuCoV-229E in MRC-5 lung cells. Our cells were maintained in culture medium supplemented with 1 mM GlutaMax-1 (L-alanyl-L-glutamine); theirs with 1 mM L-glutamine which is known to bind to copper and its spontaneous break down at physiological pH releasing ammonia during cell culture also reacts with copper to precipitate Cu(OH)_2_. These two chemistries would give a partial passivation effect, making the copper surfaces less antiviral while GlutaMAX™-1-fed viral cultures would not passivate the copper; hence explaining the longer time of glutamine propagated virus for copper inactivation. Indeed, GlutaMAX-1 has superseded glutamine for cell culture studies because of its greater stability and we have also used it as a supplement for propagating norovirus (7).

Nevertheless, both our work with Hu-CoV-229E and SARS-CoV-2, and that of van Doremalen *et al*. (5), show copper alloy surfaces are better than other surfaces for inactivating coronaviruses and preventing their spread via fomite contact. Our many laboratory studies describing the antibacterial, antiviral and antifungal properties of copper alloys have now been successfully translated into healthcare settings where worldwide studies have shown a 90% reduction in bioburden on touch surfaces in hospitals and greater than 58% reduction in infection rates in intensive care units and paediatric units (10, 12).

Although copper alloys and impregnated fabrics are being deployed worldwide in healthcare and public transportation systems, it surprises many that they have yet to find even greater acceptance as a self-disinfecting surface. It has been speculated that, despite evidence of the ability of copper to reduce microbial surface contamination, the cost of replacement of existing commonly touched objects (door handles, push plates, hand rails etc) makes it prohibitively expensive. In fact, the payback time for a hospital installing copper alloy touch surfaces, by reducing mortality and morbidity rates, is only several months (10). This can be improved even further by considering using robust copper coatings, for example using the high pressure sprayed copper coating used in this study. The process involves the acceleration of powdered copper to hypersonic speeds, using compressed gas and De LaVal nozzles. The accelerated copper particles then impact on the surface to be coated and bond, due to plastic deformation, resulting in a coating with similar adhesive strength as a traditional weld. We investigated a deposited copper layer of approximately 150 microns in depth and this proved as antiviral as pure copper. As such, it is a simple, rapid and cheap process to coat existing metals and even plastic surfaces that might be welcome in money conscious industries or low income countries.

It might be too late for deploying copper in the current COVID-19 pandemic but is notable that despite the superb efforts to rapidly develop vaccines against SARS-CoV-2 infection, beating all records, it has taken almost 12 months for the first approved vaccines to be deployed. Installing potent antimicrobial copper alloys and coatings into healthcare and public transport systems offers a simple defence in the fight to delay the spread of future pandemic diseases until vaccines become available.

